# Precise 3D Localization of Intracerebral Implants with a simple Brain Clearing Method

**DOI:** 10.1101/2023.12.22.573088

**Authors:** Julien Catanese, Tatsuya Murakami, Paul J. Kenny, Ines Ibanez-Tallon

## Abstract

Determining the localization of intracerebral implants in rodent brain stands as a critical final step in most physiological and behaviroral studies, especially when targeting deep brain nuclei. Conventional histological approaches, reliant on manual estimation through sectioning and slice examination, are error-prone, potentially complicating data interpretation.

Leveraging recent advances in tissue-clearing techniques and light-sheet fluorescence microscopy, we introduce a method enabling virtual brain slicing in any orientation, offering precise implant localization without the limitations of traditional tissue sectioning.

To illustrate the method’s utility, we present findings from the implantation of linear silicon probes into the midbrain interpeduncular nucleus (IPN) of anesthetized transgenic mice expressing chanelrhodopsin-2 and enhanced yellow fluorescent protein under the choline acetyltransferase (ChAT) promoter/enhancer regions (ChAT-Chr2-EYFP mice). Utilizing a fluorescent dye applied to the electrode surface, we visualized both the targeted area and the precise localization, enabling enhanced inter-subject comparisons. Three dimensional (3D) brain renderings, presented effortlessly in video format across various orientations, showcase the versatility of this approach.

## 1 Introduction

The field of neuroscience has experienced significant progress in recent years, propelled in large part by the utilization of intracerebral implants in *in vivo* rodent studies. Indeed, discipline such as *in vivo* electrophysiology, optogenetics, and fiber photometry all rely on intracerebral implants. Central to the validity of these studies is the histological verification of implant placement, which is crucial for the accurate interpretation of experimental outcomes. Various strategies have emerged within in vivo electrophysiology to enhance the histological identification of electrode implants, including micro-lesions employing electrolytic (Summerlee et al., 1982), radiofrequency (Brozoski et al., 2006), or heat (Chen & Wise, 1997) techniques. However, these methods, which rely on repetitive manual procedures, are time-consuming and prone to potential damage or loss of tissue slices.

Newer approaches aim to preserve the entire brain, such as brain-wide imaging of probe tracks using lightsheet microscopy and serial block-face two-photon (SBF2P) microscopy (Liu et al., 2021), a revolutionary development for histology.

In this article, we propose an optimized version of the method introduced by Liu and colleagues (2021). The previous protocol described an alcohol-based brain clearing procedure, which requires long delipidation up to 14 days, and the compatibility with fluorescent protein was not fully demonstrated (Liu et al. (2021). Our technique involves using CUBIC to clear the brain, coupled with an easy-to-follow recipe suitable for any laboratory equipped with a light-sheet microscope. Our clearing method is not only faster and safer, but it also allows for parallel processing, significantly enhancing productivity. Moreover, it is compatible with fluorescent proteins, opening new avenues for research strategies.

We introduce a novel and simple methodology for the 3D reconstruction of various implant types by simply coating them with a fluorescent dye, DiI, eliminating the need for actual brain tissue sectioning through virtual slicing. Leveraging recent breakthroughs in safe tissue clearing (Matsumoto et al., 2019b; Murakami et al., 2018; Tainaka et al., 2018b) combined with fluorescent light-sheet microscopy (Gao et al., 2015), our approach involves computational processing of virtual slices of a whole-mouse brain to reveal the exact position of implants alongside additional fluorescent tags like GFP or mCherry.

Moreover, recent advancements in image registration facilitate the automatic tracing of implant paths across cerebral structures This protocol is safe, cost-effective, and allows for the preservation of tissue slices in digital media. Our methodology, while an initial step, holds promise for further refinement and potential establishment of new standards in histological reconstruction and determination of implant location.

To illustrate the utility of this new methodology, we present experimental results involving acute implantation of both a silicon probe and an optical fiber in a deep brain area in anesthetized transgenic mice. Specifically, the electrodes were surgically placed within the Interpeduncular Nucleus (IPN), a small nucleus within the ventral midbrain known for its intricate targeting, necessitating precise angular implantation, and resulting in variability in electrode paths across subjects. Utilizing EYFP fluorescence from habenular cholinergic terminals expressing ChAT-Chr2-EYFP in transgenic mice, which densely and selectively innervate the IPN, we reliably identified the IPN in whole-brain preparations. This enabled an accurate comparison of electrode positions relative to the fluorescent IPN in the videos. Finally, we demonstrate the process of 3D electrode placement reconstruction using freely available tools from existing literature.

## 2 Methods

### 2.1 Animals and Surgeries

Experimental subjects comprised ChAT-ChR2-EYFP BAC transgenic mice (JAX stock #014546; Zao et al., 2011), adult males and females aged between 2 and 6 months, weighing 25–35g. Mice were deeply anesthetized with isoflurane (4% initiation) and maintained under continuous anesthesia (1%) throughout the procedure using a digital stereotaxic apparatus (Kopf). An articulating stereomicroscope facilitated precise craniotomy and acute electrode array implantation. Upon completion of the electrophysiological recording, mice were euthanized via Nembutal overdose and subsequently perfused intracardially with 1×PBS followed by PFA (4%).

All procedures involving mice were approved by The Rockefeller University Institutional Animal Care and Use Committee (IACUC) and were in accordance with National Institutes of Health guidelines.

### 2.2 Electrodes and implantation procedure

We employed silicon micro-machined electrode arrays, a cutting-edge tool offering several advantages over traditional insulated metal electrodes and tetrodes. These arrays provide enhanced recording capabilities, rigidity, reliability, and customizability, making them a precise choice for electrophysiological recordings.

The implanted device comprised an optrode, consisting of a linearly arranged 32-channel silicon probe (A1x32-7mm-25-177, Neuronexus) and a 100 μm diameter optic fiber mounted on the shank, terminating 150 μm from the first channel and 975 μm from the tip. The shank measured 7 mm in length, 50 μm in thickness, and 145 μm in width until the first channel (located 700 μm above the tip). The width linearly decreased from the first to the last channel, reaching 27 μm at the tip. Recording sites were spaced linearly by 25 μm, each covering a 177 μm^2^ circular surface area, with an internal reference (4200 μm^2^) positioned 50 μm above the upper channel.

Prior to implantation, the shank and tip of the silicon probe were meticulously painted or dipped into a far-red fluorescent DiO dye. This step ensured accurate visualization without compromising brain tissue.

Following careful anesthesia of the Chat mouse, we acutely implanted the silicon probe deep into the midbrain, targeting the IPN region. This procedure was executed with precision, leveraging stereotaxic ultra-precise Z movement in an entirely immobilized subject.

### 2.3 Sample Preparation steps and CUBIC Preparation

Upon completing the experiment and intracardiac perfusion, the entire brain was meticulously extracted and immersed in a 4% PFA fixative solution overnight at 4°C. Subsequently, it was transferred to a 30% sucrose solution at 4°C, prepared for the clearing procedure.

CUBIC-L was prepared by mixing 10wt% N-butyl-diethanolamine and 10wt% Triton X-100 in H_2_O, stirred, and stored at room temperature.

CUBIC-R was prepared by combining 45wt% antipyrine and 30wt% nicotinamide in H_2_O. The solution underwent brief heating (1 min at maximum microwave power) until dissolved (40∼80 sec) and was stirred at room temperature until entirely clear. It was then stored at room temperature while protected from light.

The refractive index (RI) was verified to be RI = 1.525.

A mixed oil solution for immersion (RI = 1.525) was prepared mixing silicon oil, phenylmethylsiloxane oligomer and mineral oil at the ratio of 0.6517:0.3483, stirred, and stored at room temperature.

### 2.4 Clearing Procedure details and timeline

Tissues underwent clearing utilizing the CUBIC protocol (Matsumoto et al., 2019a; Murakami et al., 2018; Tainaka et al., 2018a), obviating the need for toxic chemicals. The rapid clearing method described by Matsumoto et al., 2019, designed for highly transparent mouse organs, was chosen to facilitate the detection of electrode placement in the brain. The procedure can be summarized in three main steps:

#### 1. PBS Washes to Remove Fixative

Full brain samples were gently washed for 2 hours in 1×PBS with gentle agitation at room temperature, shielded from light. This step was repeated four times.

An optional step included reducing brain volume to expedite the process. For instance, in our case, an acrylic mouse brain matrix was utilized to carefully cut 3 mm from each extremity (on the AP axis) with a small blade, away from the region of interest.

#### 2. CUBIC-L for Delipidating Brains

Brains were immersed in 5 ml of CUBIC-L solution with gentle shaking at 37°C for 2 days.

The CUBIC-L solution was refreshed every 2 days until the brain became translucent white, around days 5-6 in our case, though this duration may vary based on mouse age and fixation procedure.

#### 3. CUBIC-R for Refractive Index (RI) Matching

Delipidated brains were immersed in 5 ml of half-diluted CUBIC-R overnight at room temperature, shielded from light with gentle agitation.

Subsequently, the brain was immersed in 4 ml of CUBIC-R in a 5-ml tube, gently shaken at room temperature, and incubated for an additional 24 hours. This step was repeated twice.

Post-clearing, brains were immediately imaged. However, they can be stored at room temperature in the dark for an extended period before imaging. Additional staining steps or pauses can be incorporated into this procedure as outlined in Matsumoto et al., 2019.

### 2.5 Light-Sheet Imaging setup and parameters

The Bio-Imaging Resource Center at The Rockefeller University provided light-sheet microscopes for imaging cleared brain samples. We employed the Ultramicroscope II (LaVision BioTec) along with ImspectorPro software (LaVision BioTec). Illumination was generated using laser sources set at appropriate wavelengths (445, 488, 561, 640, and 785 nm Lasers, LaVision BioTec).

Brain samples were placed in a 100% quartz imaging cuvette (LaVision BioTec) filled with mixed oil. The positioning of the brain defined the orientation of the virtual slices, including coronal, sagittal, horizontal, or other orientations. The illumination scanned the entire brain from top to bottom (Z-scan) with a light sheet emitted from the sides via laser light through two parallel cylindrical lenses.

For imaging, we mounted a 1.3× objective lens on a microscope stand with corrected optics (1.3×/0.1 LVMI-Fluar objective with 9 mm WD), using 1× zoom magnification to receive the emitted light from the sample’s top. Images were captured using an Andor Neo sCMOS camera (5 μm x 5 μm pixel size).

To expedite acquisition (approximately 15 minutes/brain), we utilized specific parameters:

Light sheet thickness: 6.24 μm (NA in system software set to 0.041) Unilateral acquisition with a 3 μm step size in Z-orientation

Unidirectional light sheet acquisition speed: 400 ms per image, resulting in imaging 1500-2000 slices per brain

Width of the light sheet: 60%

Laser power: set between 60-90%

Emission filters: 525/50 nm for 488 nm excitation; 680/30 nm for 640 nm excitation Image Processing and Video generation

Fluorescence images in Z-series (in .tiff format) obtained from Ultramicroscope II were processed using the Fiji software (Schindelin et al., 2012, 2015). Color assignments were arbitrary; here, we assigned green to GFP (in ChAT expressing habenular cell body and terminals within the IPN) and red to the electrode dye. Movies (.avi) were generated using Fiji at 100 frames per second (refer to video1 and video2).

### 2.6 Registration

To localize and reconstruct the implant’s path within the brain, we employed SHARP-Track, a tool developed by Shamash et al., 2018. SHARP-Track provides a MATLAB user interface to register virtual slice images relative to the Allen Mouse Brain Atlas (Lein et al., 2006; Oh et al., 2014; Sunkin et al., 2013).

## 3 Results

### 3.1 Overview of the Procedure for Precise Electrode Localization and Brain Visualization

The procedure’s primary goal was to achieve a rapid and accurate readout of electrode positioning within the fully cleared brain while visualizing areas expressing fluorescent proteins of interest. Employing the clearing procedure in conjunction with light sheet microscopy allowed for comprehensive brain visualization in various orientations without the need for sectioning. The procedural steps (see Figure 1) were straightforward, expeditious, and facilitated the simultaneous processing of multiple brain samples, thus optimizing research time and mitigating the inherent risks of manual labor-related errors.

**Figure 1:**
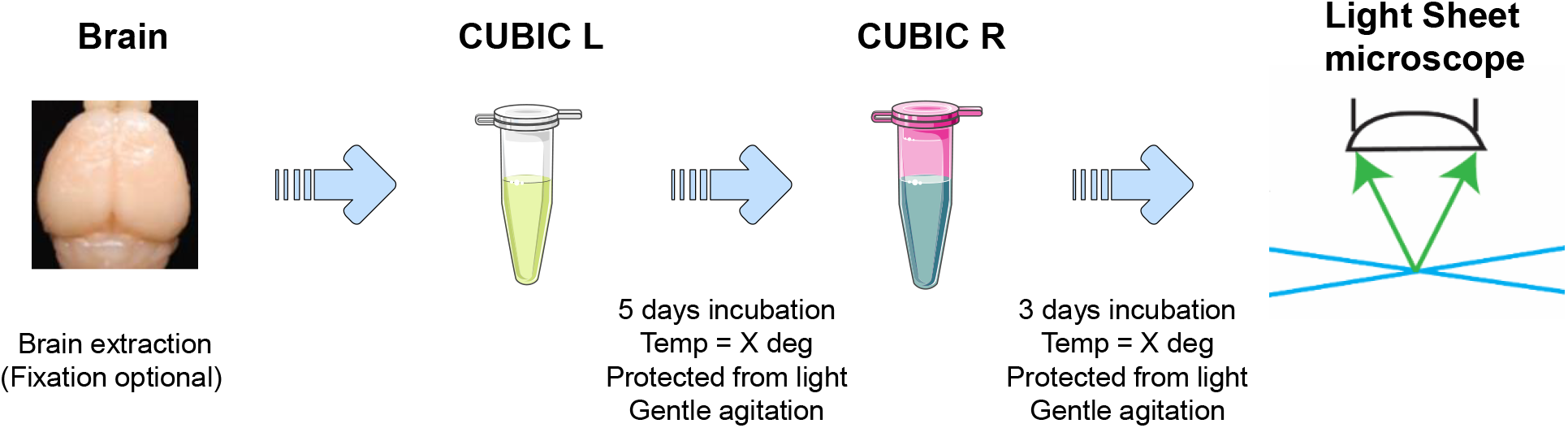
Procedure Schematic. A Schematic Overview of the Procedure. We employed silicon probes for acute recordings in the Interpeduncular Nucleus (IPN) of Chat-CHR2-EYFP mice under anesthesia. Post-experiment brain processing involved minimal manual labor. The clearing procedure necessitated only periodic reagent changes and refreshments every 1-2 days (refer to methods).

#### Procedure Summary

- **Brain Preparation:** Brains were fixed and extracted within 1-24 hours post in-vivo experiments, stored in sucrose solution at 4°C for 1 to 90 days, and subsequently washed with PBS.
- **Delipidation and Refractive Index Adjustment:** Brain samples underwent delipidation by incubating in CUBIC-L solution at 37°C for 5-6 days, followed by washing with PBS. Subsequently, brains were incubated in CUBIC-R solution for 2-3 days to achieve the desired refractive index.
- **Imaging Procedure:** Using a light-sheet fluorescent microscope, brain samples were imaged with virtual sectioning, obtaining 2D images every 3 μm within 5-10 minutes for 1000-2000 tiff images/sample. Imaging was conducted with different excitation/emission wavelengths and various virtual section plans (e.g., coronal and sagittal).
- **Video Generation and Analysis:** Image processing through FIJI (ImageJ) facilitated quick conversion of brain images into colored videos. The processing time for video creation was 10-20 minutes with a 32G RAM computer. Videos enabled rapid visualization of electrode localization and fluorescent protein expression in the region of interest (Refer to Figures 2 and 3). These videos were easy to store and share, simplifying data interpretation.

**Figure 2:**
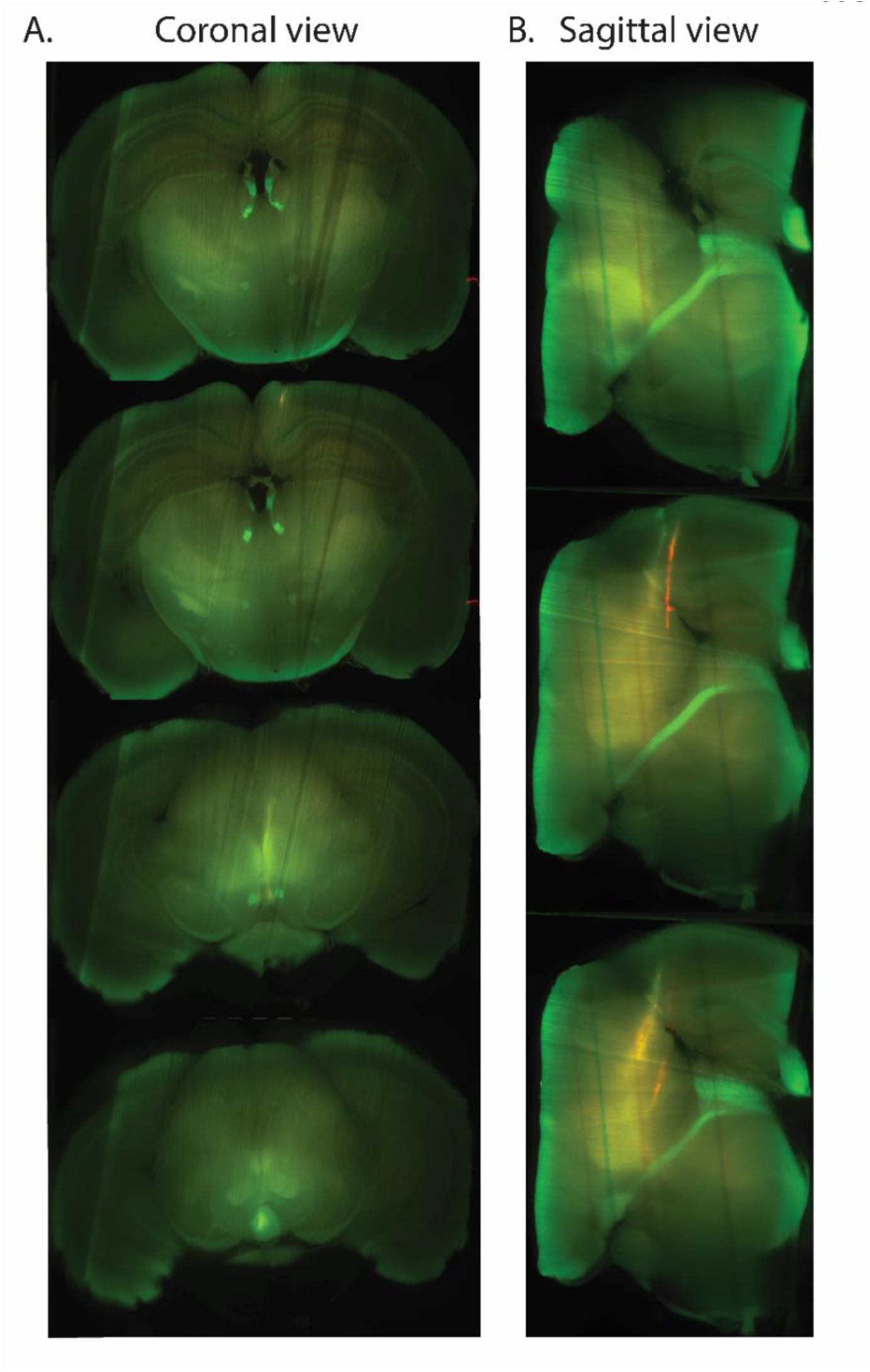
Revealing Fluorescent Pathways Across Various Planes. **A. Coronal View:** Four virtual slices obtained from a mouse cleared brain sample using the light sheet microscope. Bright green areas depict cholinergic neurons highlighting the mHb in the top slice. The Fasciculus Retroflexus (FR) can be traced until its convergence into the IPN (bottom slice). **B. Sagittal View:** Three virtual slices from the same brain as in A. The elongated bright green path observed from the side represents the FR extending from the mHb to the IPN. Additionally, the red color indicates a dye (DiO) applied to the shanks of the silicon probes. In both panels (A and B), note the presence of straight lines traversing the image, indicative of noise from the light sheet microscope. These artifacts can be distinguished from the objects of interest, namely, electrodes (red) and cholinergic tissue (green).

**Figure 3:**
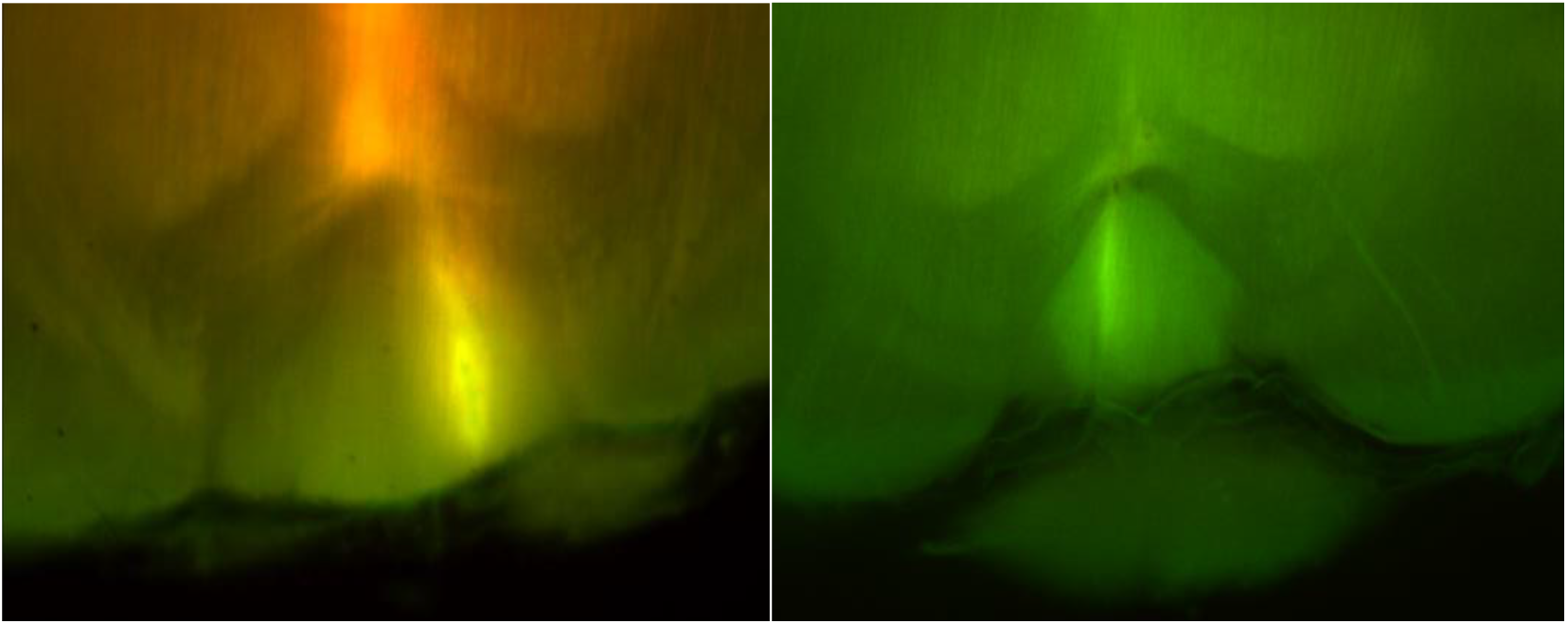
Example of Silicon probe tip visualization within the IPN. The two pictures are from two independent experiments, and both zoomed on the IPN region. The IPN (green) appear in green. The silicon probe tip (red) appears in yellow as the result of the merging of the red image on the green image. IPN.

### 3.2 Revealing Fluorescent Brain Circuits

Figure 2 illustrates the successful visualization of the targeted brain circuit through our procedure. In this experiment, our aim was to observe the mHb-IPN pathway. To achieve this, we utilized transgenic mice expressing Yellow Fluorescent Protein (YFP) under the Choline Acetyl Transferase (ChAT) promoter, specific to cholinergic neurons involved in Acetylcholine synthesis. The dense presence of cholinergic neurons in the medial Habenula (mHb), projecting directly to the Interpeduncular nucleus (IPN), is prominently visible in Figure 2 and the supplementary videos. Although the signal-to-noise ratio isn’t perfect, it is adequate to observe the pathway of interest. Therefore, we opted not to further improve this aspect. In conclusion, our clearing methods effectively revealed the specific mHb-IPN pathway, as showcased in Figure 2 and the accompanying videos.

One of the significant advantages of this technique lies in its ability to perform virtual brain sectioning across various dimensions or plans of virtual cut. Figure 2 demonstrates the identical brain under two different classical dimensions: coronal and sagittal. In this illustration, the Fasciculus Retroflexus (FR) can be distinctly identified in the Sagittal view as a lengthy bundle of axons originating from the medial Habenula (mHb) and traversing through the entire brain, reaching deep into the Interpeduncular Nucleus (IPN). Conversely, in a coronal view, two spots of light (left and right bundle) gradually descend ventrally until merging within the IPN. The capacity to visualize the brain from multiple angles enables more flexible tracking of intracerebral objects of interest.

### 3.3 Revealing Electrode Tracks Localization within Brain Circuit

The primary objective of precisely mapping the electrode’s location within the cleared brain has been effectively achieved has shown in Figure 3.

Figure 3 vividly displays the culmination of our approach, presenting a striking image where the red fluorescence indicative of the dye (DiD, far red) from the electrode tip, precisely nestled within the green fluorescence denoting the Interpeduncular Nucleus (IPN) marked by GFP expression. This remarkable alignment provides a clear visual depiction of our achievement in precisely pinpointing the electrode location within the intended brain region.

### 3.4 Registration for Precise 3D Cross-Comparison

The registration process facilitates cross-comparison between brains from various experiments and subjects, addressing inter-individual variability. While numerous experiments generate distinct brain image sets, aligning these data into a common coordinate system enables accurate comparisons. In our approach, we opted for the Allen Brain Atlas due to its continuous 3D representation, unlike the Paxinos Atlas, which is more suitable for brain slice data registration. Despite the absence of a dedicated registration tool in the Allen Brain Atlas, various groups have developed their solutions (Bakker et al., 2015; Puchades et al., 2019). However, a universally accessible, straightforward solution remains elusive.

Our proposal utilizes the SHARP-Track tool (developed by Shamash et al., 2018), offering a simple and freely available method, as depicted in Figure 4, yielding satisfactory results to precisely localize the brain areas crossed by the probe shank. While alternative programs could achieve similar outcomes, we selected this tool due to its user-friendly interface. Moreover, our imaging method harmonizes seamlessly with SHARP-Track, generating virtual slices across various dimensions. Despite the visual inference of electrode locations via images (Figure 3), employing the registration steps enhances the identification of brain subregions intersected by the electrode path (Figure 4).

**Figure 4:**
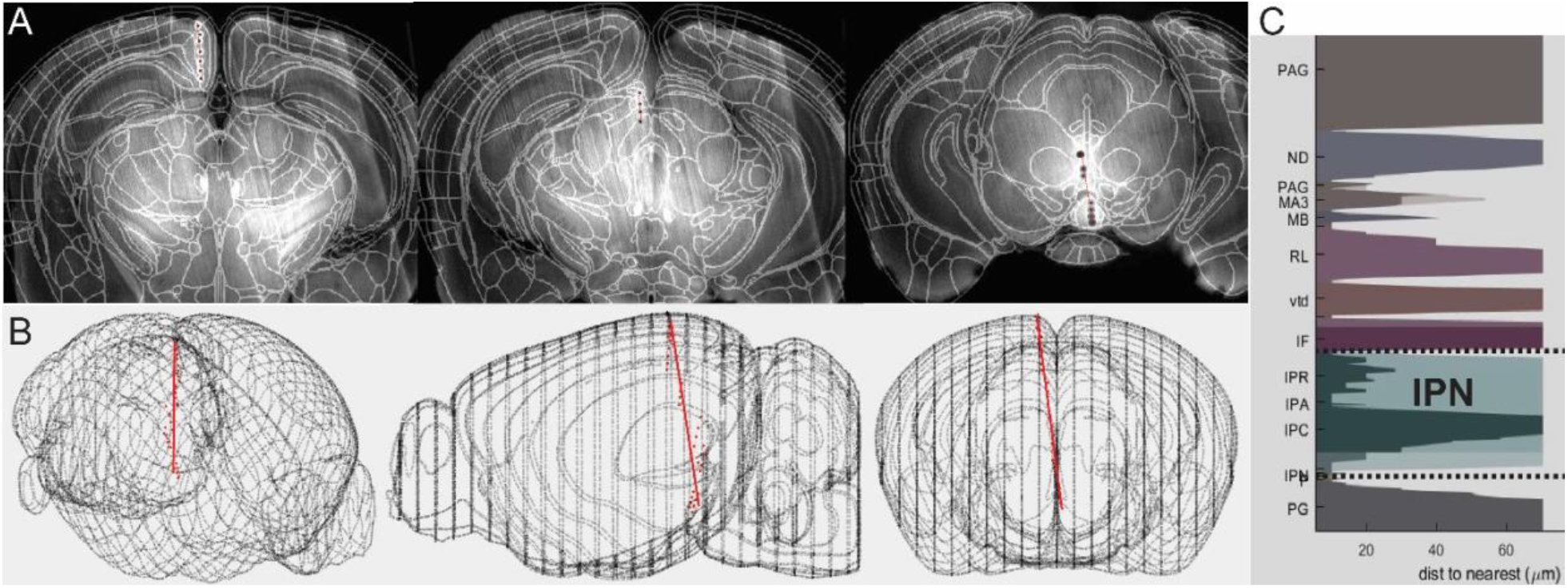
Registration and Identification of Brain Regions Traversed by a Probe. **(A) Brain images**, sliced virtually in any direction using the light-sheet microscope, were registered to the reference atlas. This process was executed with SHARP-Track, a free software developed by Shamash et al., 2018. The electrodes were manually identified, marked by black dots along the red line. **(B) Utilizing the SHARP-Track tool**, best-fit lines for the electrodes were computed in 3D within the mouse brain. **(C) identifying brain regions traversed by the probe track**. Brain region identities are displayed on the y-axis, represented by acronyms from the Allen Brain Atlas, utilized for registration. The thickness along the x-axis provides an estimation of the confidence level in the accuracy of each brain-region label (see Shamash et al., 2018 for details). Arbitrary colors are used to denote regions belonging to the same parent region. As anticipated, all the IPN sub-regions (IPR, IPA, IPC, and IPNP) are identified in green. Dotted lines delineate the extent of the electrode’s recording channels.

### 3.5 Conclusion

Our experimental methodology has proven effective in localizing and visualizing electrodes within the brain’s circuitry. The success of our approach confirms its accuracy in precisely identifying and tracking electrode placement within the complex neural networks of the brain.

This visualization underscores the precision of our technique, emphasizing its potential for advancing neuroscientific investigations. By successfully navigating the intricate neural landscape, our study contributes valuable insights into understanding the functioning of the brain.

In addition to its significance in basic research, the practical applications of our methodology extend to clinical settings. The precise identification of electrode placement holds promise for the development of neurostimulation therapies and brain-machine interfaces.

In summary, our experimental results demonstrate the effectiveness of our methodology in localizing and visualizing electrodes within the brain. This achievement has implications for both basic research and potential clinical applications, offering a clearer understanding of the brain’s intricate neural networks.

## 4 Discussion and Limitations

### 4.1 Bridging Technology for Enhanced Brain Mapping

Our novel method combines cutting-edge brain clearing protocols (CUBIC) and light-sheet microscopy to precisely identify electrode locations alongside fluorescent molecules in transgenic mouse brains. This innovative approach significantly outperforms conventional histological techniques, offering heightened accuracy, reduced data loss, and streamlined execution, even for individuals with limited expertise in the field.

### 4.2 A Timeline for Parallel Processing

Upon adopting this new clearing method over classical histology, we discovered its significant advantages in time management and its potential to enhance productivity. To illustrate, here is the timeline we implemented for our experiments.

Each week, we conducted operations and recorded data from 3-4 mice (1 mouse per day). At the end of each day, brains were fixed and kept at 4°C. Processing of brains can be parallelized during each cycle (we suggest a 3-week cycle). With the clearing steps allowing for hands-free work, it becomes feasible to conduct experiments simultaneously, enabling overlapping cycles (initiating a new cycle each week). This approach maintains a continuous flow of data acquisition, and the entire process—from experiments to analysis—can be managed by a single individual following this timeline.

### 4.3 Methodological Preferences and Technological Advancements

Among various available clearing methods, our preference for the CUBIC procedure stems from its enhanced safety profile and compatibility with fluorescent protein compared to alternatives like iDISCO. The utilization of light-sheet microscopy for imaging enables rapid, comprehensive scans of the entire brain, facilitating fluorescence visualization. Leveraging FIJI software further simplifies data processing, enabling the creation of accessible, informative colored videos, even for non-specialists.

### 4.4 Advancements in Neuroscientific Methodology

In the context of burgeoning interest in brain clearing technology, our method represents a pioneering approach toward advancing histological 3D reconstruction in neuroscience. While alternative techniques like micro-CT imaging with electron microscopy staining have been proposed, our unique capability to visualize fluorescent molecules, particularly in conjunction with transgenic technology, stands out for assessing neuronal brain activity.

### 4.5 Safety, Compatibility, and Future Improvements

Unlike electron microscopy staining techniques involving hazardous reagents, our approach prioritizes user safety without compromising efficacy. Although microscopic magnetic resonance imaging (μMRI) is available for brain volume reconstruction, its limitations with in vivo electrode implants and high costs restrict its practical utility in our experimental framework.

### 4.6 Expanding Methodological Scope and Anticipating Future Refinements

Our optional use of antibodies within the CUBIC procedure for immunofluorescence marking presents added versatility, albeit with a temporal trade-off due to incubation requirements. To enhance our method’s efficacy, mitigating brain tissue expansion during clearing through experimental adjustments holds promise for future refinements.

### 4.7 Methodological Adaptability and Technological Improvements

While our study demonstrates successful silicon probe localization, further validation is needed for single wire electrodes or tetrodes arrays. The optimization for acute probe implantation provides a platform for future adaptations for chronic implants. Improving image quality by refining parameters and exploring subcellular zooming capabilities could yield richer data insights.

### 4.8 Vision for Integration and Automation

Considering advancements in artificial intelligence, there’s potential for streamlined registration steps and automated processes, such as those illustrated in Figure 4. This integration could minimize human intervention, bolstering efficiency in data analysis and interpretation.

### 4.9 Pioneering Advancements in Histology

We see our method as a foundational step towards updating traditional section-based histological procedures. This could mark a shift in histology towards incorporating technological advancements, potentially improving neuroscientific investigations.

## Supporting information

supplemental video Fig2 Coronal

## ACKNOWLEDGEMENTS

The Frits and Rita Markus Bio-Imaging Resource Center (RRID:SCR_017791) at The Rockefeller University. The Heintz lab at the Rockefeller University. Grant UG3 DA048385/DA/NIDA to I.IT, Japanese Society for Promotion of Science Overseas Research Fellowship (to T.C.M)

